# Rapid Identification of Methylase Specificity (RIMS-seq) jointly identifies methylated motifs and generates shotgun sequencing of bacterial genomes

**DOI:** 10.1101/2021.03.08.434449

**Authors:** Chloé Baum, Yu-Cheng-Lin, Alexey Fomenkov, Brian P. Anton, Lixin Chen, Thomas C. Evans, Richard J Roberts, Andrew C Tolonen, Laurence Ettwiller

## Abstract

DNA methylation is widespread amongst eukaryotes and prokaryotes to modulate gene expression and confer viral resistance. 5-methylcytosine (m5C) methylation has been described in genomes of a large fraction of bacterial species as part of restriction-modification systems, each composed of a methyltransferase and cognate restriction enzyme. Methylases are site-specific and target sequences vary across organisms. High-throughput methods, such as bisulfite-sequencing can identify m5C at base resolution but require specialized library preparations and Single Molecule, Real-Time (SMRT) Sequencing usually misses m5C. Here, we present a new method called RIMS-seq (Rapid Identification of Methylase Specificity) to simultaneously sequence bacterial genomes and determine m5C methylase specificities using a simple experimental protocol that closely resembles the DNA-seq protocol for Illumina. Importantly, the resulting sequencing quality is identical to DNA-seq, enabling RIMS-seq to substitute standard sequencing of bacterial genomes. Applied to bacteria and synthetic mixed communities, RIMS-seq reveals new methylase specificities, supporting routine study of m5C methylation while sequencing new genomes.

## Introduction

DNA modifications catalysed by DNA methyltransferases are considered to be the most abundant form of epigenetic modification in genomes of both prokaryotes and eukaryotes. In prokaryotes, DNA methylation has been mainly described as part of the sequence-specific restriction modification system (RM), a bacterial immune system to resist invasion of foreign DNA (Loenen et al., 2014). As such, profiling methylation patterns gives insight into the selective pressures driving evolution of their genomes.

Around 90% of bacterial genomes contain at least one of the three common forms of DNA methylation: 5-methylcytosine (m5C), N4-methylcytosine (m4C), and N6-methyladenine (m6A))(Beaulaurier et al., 2019; Blow et al., 2016). Contrary to eukaryotes where the position of the m5C methylation is variable and subject to epigenetic states, bacterial methylations tend to be constitutively present at specific sites across the genome. These sites are defined by the methylase specificity and, in the case of RM systems, tend to be fully methylated to avoid cuts by the cognate restriction enzyme. The methylase recognition specificities typically vary from 4 to 8 nucleotides and are often, but not always, palindromic (Roberts et al., 2015).

PacBio Single Molecule, Real-Time (SMRT) sequencing has been instrumental in the identification of methylase specificity largely because, in addition to providing long read sequencing of bacterial genomes, m6A and m4C can easily be detected using the characteristic interpulse duration (IPD) of those modified bases (Flusberg et al., 2010). Thus, a single run on PacBio allows for both the sequencing and assembly of unknown bacterial genomes and the determination of m6A and m4C methylase specificities. However, because the signal associated with m5C bases is weaker than for m6A or m4C, the IPD ratio of m5C is very similar to the IPD of unmodified cytosine. Thus, PacBio sequencing usually misses the m5C methylases activities (Blow et al., 2016). Instead, the identification of m5C requires specialized methods such as bisulfite sequencing or enzyme-based techniques such as EM-seq (Sun et al., 2021). Recently, MFRE-Seq has been developed to identify m5C methylase specificities in bacteria (Anton et al., 2021). MFRE-Seq uses a modification-dependent endonuclease that generates a double-stranded DNA break at methylated m5CNNR sites, allowing the identification of m5C methylase specificities at a subset of sites. m5C in the CpG context has been identified (Simpson et al., 2017) and a signal for methylation can be observed at known methylated sites in bacteria using Nanopore sequencing (Rand et al., 2017) but so far no technique has allowed for the dual sequencing of genomes and the *de novo* determination of m5C methylase specificity for the non-CpG contexts typically found in bacteria.

Herein we describe a novel approach called RIMS-seq to simultaneously sequence bacterial genomes and globally profile m5C methylase specificity using a protocol that closely resembles the standard Illumina DNA-seq with a single, additional step. RIMS-seq shows comparable sequencing quality as DNA-seq and accurately identifies methylase specificities. Applied to characterized strains or novel isolates, RIMS-seq *de novo* identifies novel activities without the need for a reference genome and also permits the assembly of the bacterial genome at metrics comparable to standard shotgun sequencing.

## Results

### Principle of RIMS-seq

Spontaneous deamination of cytosine (C) leading to uracil (U) and of m5C leading to thymine (T) are examples of common damage found in DNA. *In vitro*, this damage is often undesirable as it can interfere with sequencing. The type of interference during sequencing depends on whether the deamination occurs on C or m5C. Uracil blocks the passage of high-fidelity polymerases typically used in library preparation protocols, preventing the amplification and sequencing of U-containing DNA fragments. Conversely, DNA harboring T derived from m5C deamination can be normally amplified, but results in C to T errors (Duncan and Miller, 1980; Fogg et al., 2002). This distinction between blocking and mutagenic damage forms the basis of the RIMS-seq method, allowing the identification of methylase specificity based on an elevated number of reads containing C to T transitions specifically at methylated sites (**Figure 1A**). For this, the DNA is subjected to a heat-alkaline treatment, inducing a level of deamination large enough to detect the m5C methylase specificity without affecting the sequencing quality.

**Figure 1 :**
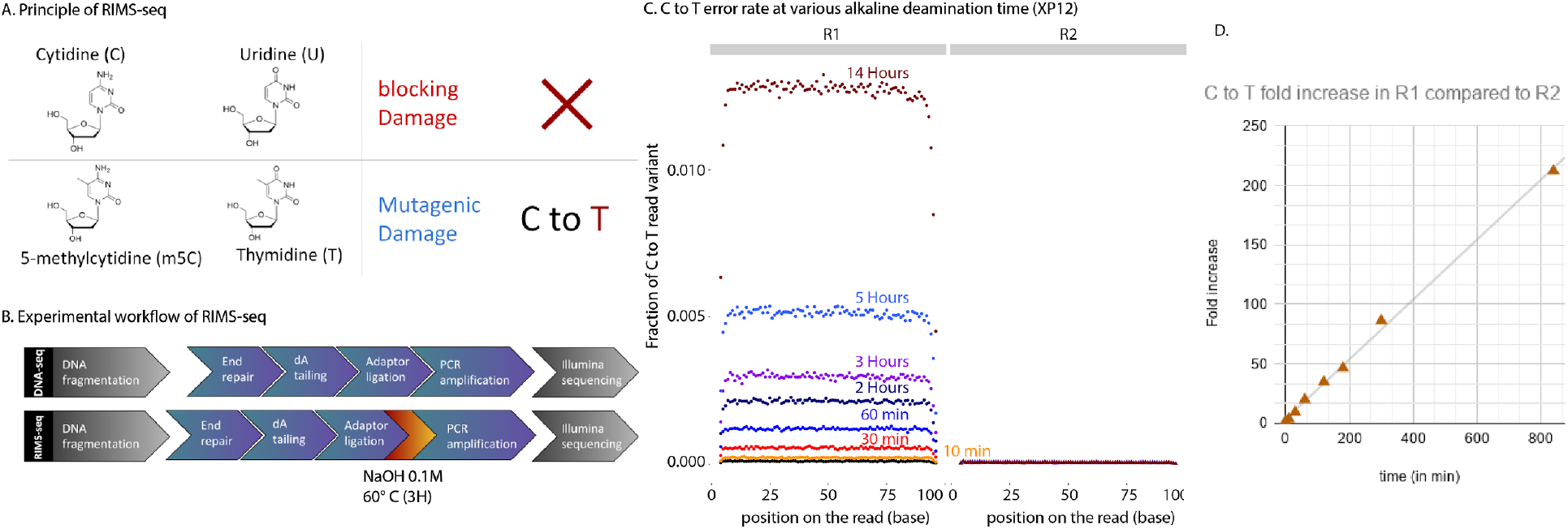
Principle of RIMS-seq. Deamination of cytidine leads to a blocking damage while deamination of m5C leads to a mutagenic C to T damage only present on the first read (R1) of paired-end reads in standard Illumina sequencing. Thus, an increase of C to T errors in R1 in specific contexts is indicative of m5C. **B.** The workflow of RIMS-seq is equivalent to a regular library preparation for Illumina DNA-seq with an extra step of limited alkaline deamination at 60°C. This step can be done immediately after adaptor ligation and does not require additional cleaning steps. **C.** Fraction of C to T variants in XP12 (m5C) at all positions in the reads for R1 and R2 after 0min (DNA-seq), 10min, 30min, 60min, 2h, 3h, 5h and 14h of heat-alkaline treatment. The C to T imbalance between R1 and R2 is indicative of deamination of m5C and increases with heat-alkaline treatment time. **D.** Correlation between the C to T fold increase in R1 compared to R2 according to time (r^2^=0.998).

Illumina paired-end sequencing allows both ends of a DNA fragment to be sequenced, generating a forward read (R1) and reverse read (R2). Resulting from m5C deamination, R1 has the C to T read variants while R2 has the reverse-complement G to A variant. This difference leads to an overall imbalance of C to T variants between R1 and R2 (Chen et al., 2017) (see also **Supplementary Figure 1** for explanation). Thus, sequence contexts for which the C to T read variants are imbalanced in R1 compared to R2 correspond to m5C methylase specificity(ies). Because of the limited deamination rate, RIMS-seq takes advantage of the collective signal at all sites to define methylase specificity. Because C to T imbalance can be observed at nucleotide resolution, RIMS-seq identifies at base resolution which of the cytosine within the motif is methylated.

The experimental steps for RIMS-seq essentially follows the standard library preparation for Illumina sequencing with an extra deamination step. Briefly, the bacterial genomic DNA is fragmented and adaptors are added to the ends of DNA fragments using ligation (**Figure 1B** and **Methods**). Between the ligation step and the amplification step, an alkaline heat treatment step is added to increase the rate of deamination. Only un-deaminated DNA or DNA containing deaminated m5C can be amplified and sequenced.

### Validation of RIMS-seq

#### Optimization of the heat alkaline deamination step

We first evaluated the conditions to maximize the deamination of m5C while minimizing other DNA damage. For this we used bacteriophage Xp12 genomic DNA that contains exclusively m5C instead of C (Kuo et al., 1968) to measure the m5C deamination rates in various contexts.

To estimate the overall deamination rate of m5C, we quantified the imbalance of C to T read variants between R1 and R2 for 0, 10 and 30 minutes, 1h, 2h, 3h, 5h and 14h of heat alkaline treatment (**Figure 1C**). We observed an imbalance as early as 10 minutes with a 3.7 fold increase of C to T read variants in R1 compared to R2. The increase is linear with time with a maximum of 212 fold increase of C to T read variants in R1 compared to R2 after 14 hours of heat alkaline treatment (**Figure 1D**). Next, we quantified the deamination rate at all Nm5CN sequence contexts with N being A,T,C or G and show an increase of C to T variants in R1 in all contexts (**Supplementary Figure 2A)**. Together, these results show that a measurable deamination rate can be achieved in as soon as 10 minutes of heat alkaline deamination and that deamination efficiency is similar in all sequence contexts.

To estimate the non-specific damage to the DNA leading to unwanted sequencing errors, we quantified possible imbalances for other variant types (**Supplementary Figure 2B**). We found that G to T variants show imbalance that is likely the result of oxidative damage resulting from sonication, a common step in library preparation between RIMS-seq and DNA-seq (Chen et al., 2017). Slight elevation of G to C and T to C read variants can be observed in RIMS-seq compared to DNA-seq, but this damage is of low frequency and therefore is not expected to notably affect the sequencing performance QC of RIMS-seq. We performed QC metrics and assemblies of Xp12 for all the alkaline-heat treatment conditions, including a control DNA-seq. The overall sequencing performances were assessed in terms of insert size, GC bias, and genome coverage. Similar results were observed between RIMS-seq and the DNA-seq control at all treatment times, indicating that the RIMS-seq heat-alkaline treatment does not affect the quality of the libraries. (**Supplementary Figure 3**).

We also evaluated the quality of the assemblies compared to the Xp12 reference genome and found that all conditions lead to a single contig corresponding to essentially the entire genome with very few mismatches (**Supplementary Table 1**). These results suggest that the heat-alkaline treatment does not affect the assembly quality, raising the possibility of using RIMS-seq for simultaneous *de novo* genome assembly and methylase specificity identification. We found that a 3 hour treatment provides a good compromise between the deamination rate (resulting in about ~ 0.3 % of m5C showing C to T transition) and duration of the experiment. We found that longer incubation times (up to 14h) increased the deamination rate by up to 1%, and decided this is a slight sensitivity increase compared to the additional experimental time required.

#### RIMS-seq is able to distinguish methylated versus unmethylated motifs in E. coli

To validate the application of RIMS-seq to bacterial genomes, we sequenced dcm- (K12) and dcm− (BL21) *E. coli* strains. In K12, the DNA cytosine methyltransferase *dcm* methylates cytosine at C<u>C</u>WGG sites (<u>C</u> = m5C, W = A or T) and is responsible for all m5C methylation in this strain (Marinus and Morris, 1973). *E. coli* BL21 has no known m5C methylation. Heat/alkaline treatments were performed at three time points (10min, 1h, and 3h). In addition, we performed a control experiment corresponding to the standard DNA-seq. Resulting libraries were paired-end sequenced using Illumina and mapped to their corresponding genomes (**Methods**).

For comparison, all datasets were down-sampled to 5 million reads corresponding to 200X coverage of the *E. coli* genome and instances of high confidence C to T variants (Q score > 35) on either R1 or R2 were identified. As expected, control DNA-seq experiments show comparable numbers of C to T read variants between R1 and R2, indicating true C to T variants or errors during amplification and sequencing (**Figure 2A**). On the other hand, the overall number of C to T read variants in R1 is progressively elevated for 10 min, 1 hour and 3 hours of heat-alkaline treatment of the *E. coli* K12 samples with an overall 4 fold increase after 3 hours treatment compared to no treatment; heat-alkaline treatments did not increase the rate of C to T read variants in R2 (**Figure 2A**). We anticipate that the elevation of the *E.coli* K12 C to T read variants in R1 is due to deamination of m5C. In this case, the elevation should be specifically found in Cs in the context of C<u>C</u>WGG (with the underlined C corresponding to the base under consideration). To demonstrate this, we calculated the fraction of C to T read variants in C<u>C</u>WGG compared to other contexts. We observed a large elevation of the C to T read variants in the C<u>C</u>AGG and C<u>C</u>TGG contexts for K12 (**Figure 2B**). As expected, the C to T read variants show no elevation at C<u>C</u>AGG and C<u>C</u>TGG contexts for the *E.coli* BL21 strain that is missing the *dcm* methylase gene (**Figure 2B**). Thus, this C to T read variant elevation is specific to the *E.coli* K12 strain subjected to heat-alkaline treatments, consistent with deamination detectable only on methylated sites. Taken together, these results indicate that the elevated rate of C to T variants observed in R1 from *E.coli* K12 is the result of m5C deamination in the C<u>C</u>WGG context. Next, we assessed whether the difference in the C to T read variant context between R1 and R2 at the C<u>C</u>WGG motif provides a strong enough signal to be discernible over the background noise. For this, we calculated the fraction of C to T read variants in C<u>C</u>WGG and C<u>C</u>WGG compared to all the other N<u>C</u>NNN and <u>C</u>NNNN contexts, respectively. After 3 hours of heat-alkaline treatment, the fraction of C to T read variants in a C<u>C</u>WGG context increased, rising from only 1.9 % in regular DNA-seq to ~25% of all the C to T variants. This increase is only observable in R1 of the K12 strain (**Figure 2C**). Conversely, no increase can be observed in <u>C</u>CWGG context for which the C to T variant rate at the first C is assessed (**Figure 2C**). Thus, RIMS-seq identified the second C as the one bearing the methylation, consistent with the well described dcm methylation of *E.coli* K12 (Palmer and Marinus, 1994) (Marinus and Morris, 1973), highlighting the ability of RIMS-seq to identify m5C methylation at base resolution within the methylated motif.

**Figure 2.**
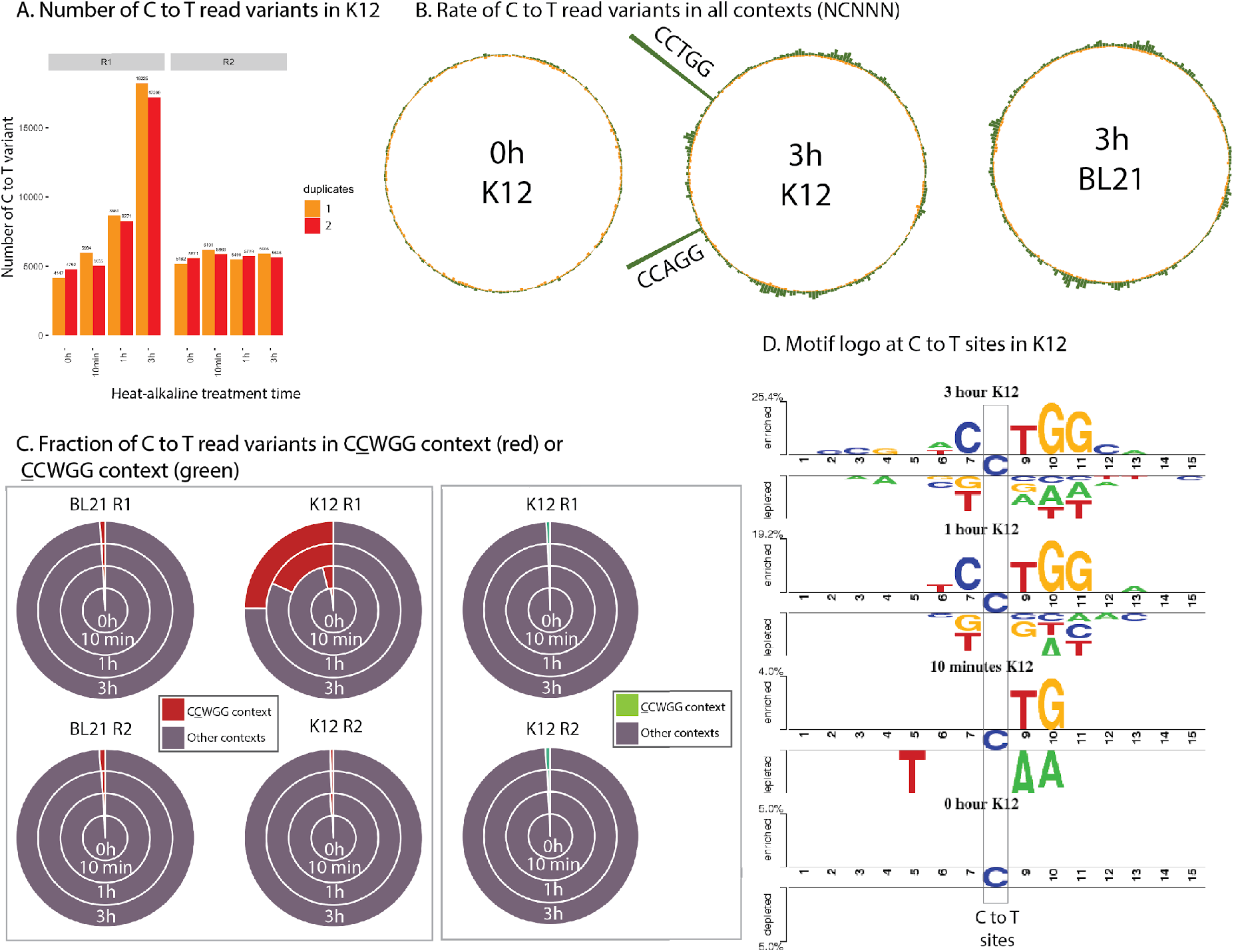
**A.** Barplots representing the number of C to T read variants for K12 in R1 and R2 after different heat/alkaline treatment times. Colors represent duplicate experiments. **B.** Circular barplot representing the rate of C to T read variants in all N<u>C</u>NNN contexts (with N = A,T C or G) for R1 (green) and R2 (orange) for DNA-seq (left) and for 3 hours heat/alkaline treatment (RIMS-seq) for K12 (center) and BL21 strains (right). **C.** Proportion of C to T read variants in C<u>C</u>WGG (red) or C<u>C</u>WGG (green) contexts compared to other N<u>C</u>NNN or <u>C</u>NNNN contexts for R1 and R2 in K12 and BL21. The C to T read variants in C<u>C</u>WGG and C<u>C</u>WGG motifs represent less than 2% of all variants except in K12 (R1 only) after 10 minutes, 1 and 3 hours treatments where the C<u>C</u>WGG motifs represent 4.1%, 22.5% and 32.6% of all C to T read variants respectively. The increase of C to T read variants in the C<u>C</u>WGG context is therefore specific to R1 in K12 strain. **D.** Visualization of the statistically significant differences in position-specific nucleotide compositions around C to T variants in R1 compared to R2 using Two Sample Logo (Crooks et al., 2004) for the K12 sample subjected to (from top to bottom) 3H, 1H, 10 min and 0 min heat alkaline treatment.

Next, we calculated significant (p.value < 0.01) differences in position-specific nucleotide compositions around C to T variants in R1 compared to R2 using Two Sample Logo (Crooks et al., 2004). We found a signal consistent with the dcm methylase specificity in K12 RIMS-seq samples at one and three hours of heat alkaline treatment (**Figure 2D**) demonstrating that it is possible to identify methylase specificities in genomic sequence subject to as little as 1h of alkaline treatment. These results support the application of RIMS-seq for the *de novo* identification of methylase specificity at base resolution.

#### RIMS-seq identifies the correct methylase specificity *de novo* in *E. coli* K12

In order to identify methylase specificities *de novo* in RIMS-seq sequencing data, we devised an analysis pipeline based on MoSDi (Marschall and Rahmann, 2009) to extract sequence motif(s) with an over-representation of C to T transitions in R1 reads (**Figure 3A**, analysis pipeline available at https://github.com/Ettwiller/RIMS-seq). In brief, the pipeline extracts the sequence context at each C to T read variant in R1 (foreground) and R2 (background). MoSDi identifies the highest over-represented motif in the foreground sequences compared to the background sequences. To accommodate the presence of multiple methylases in the same host, the first motif is subsequently masked in both the foreground and background sequences and the pipeline is run again to find the second highest over-represented motif and so on until no significant motifs can be found (see **Methods** for details). Running the pipeline on strain K12 identifies one significant over-represented motif corresponding to the C<u>C</u>WGG motif (p-value = 4.49e^−96^ and 1.10e^−1575^ for 10 min and 60 min of alkaline treatment respectively) with the cytosine at position 2 being m5C.

**Figure 3 :**
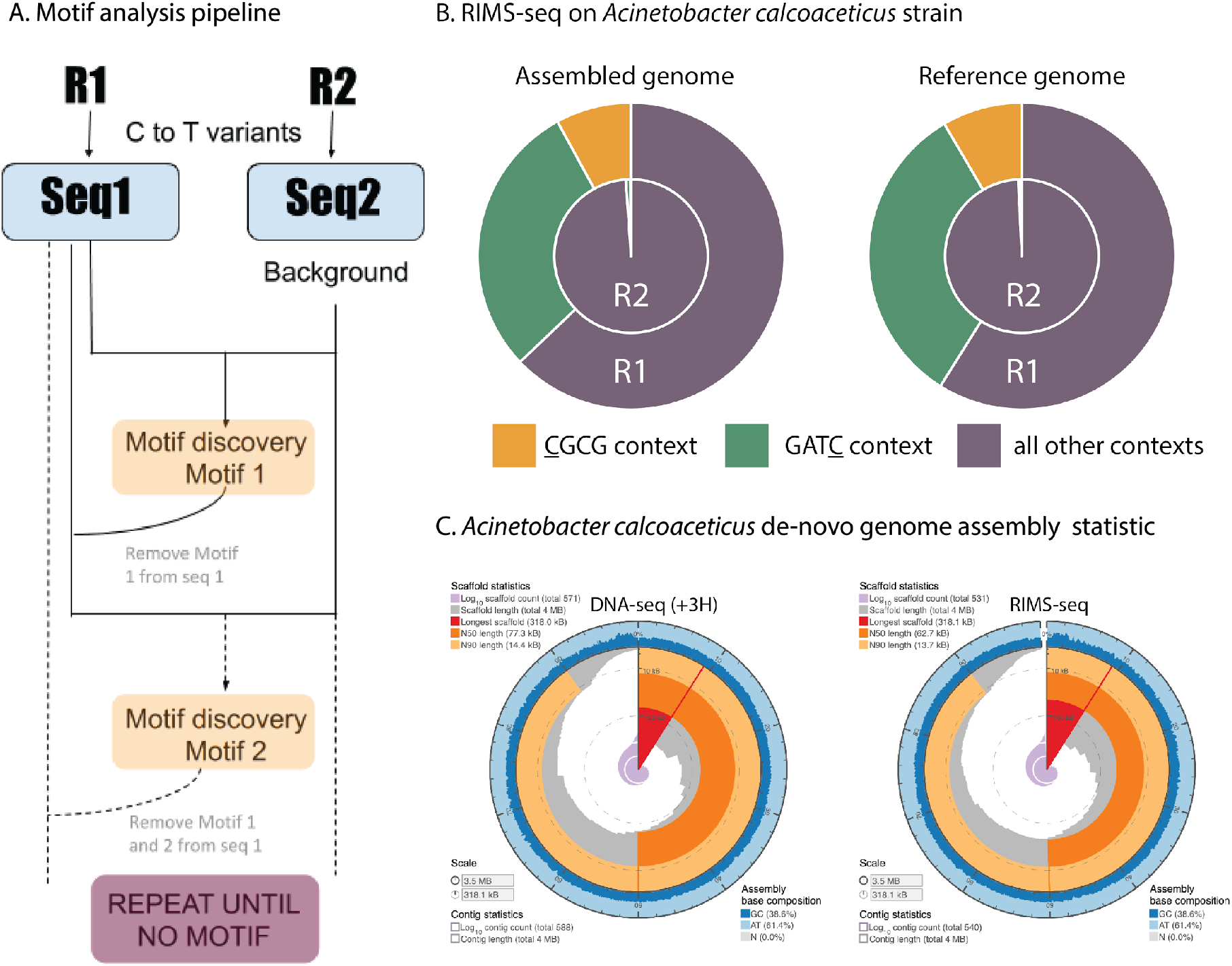
*De novo* discovery of methylase specificity using RIMS-seq. **A.** Description of the RIMS-seq motif analysis pipeline. First, C to T read variants are identified in both Read 1 and Read 2 separately. Then, the MosDI program searches for overrepresented motifs. Once a motif is found, the pipeline is repeated until no more motifs are found, enabling identification of multiple methylase specificities in an organism. **B**. Fractions of C to T read variants in <u>C</u>GCG (yellow) or GAT<u>C</u> (green) contexts compared to other contexts for R1 and R2 in *Acinetobacter calcoaceticus* ATCC 49823 using the assembled or the reference genome. The increase of C to T read variants in these contexts is similar when using either the assembled or reference genomes **C**. Assembly statistics obtained using the sequence from the standard DNA-seq (+3H, left) and RIMS-seq (right). Visualization using assembly-stats program (https://github.com/rjchallis/assembly-stats). The corresponding table with the statistical values is available in the supplementary material (**Supplementary Table 2**).

Summing up, we devised a novel sequencing strategy called RIMS-seq and its analysis pipeline to identify m5C methylase specificity *de novo*. When applied to *E. coli* K12, RIMS-seq identifies the dcm methylase specificity as C<u>C</u>WGG with the methylated site located on the second C, consistent with the reported dcm methylase specificity (**Table 1**).

**Table 1 :**
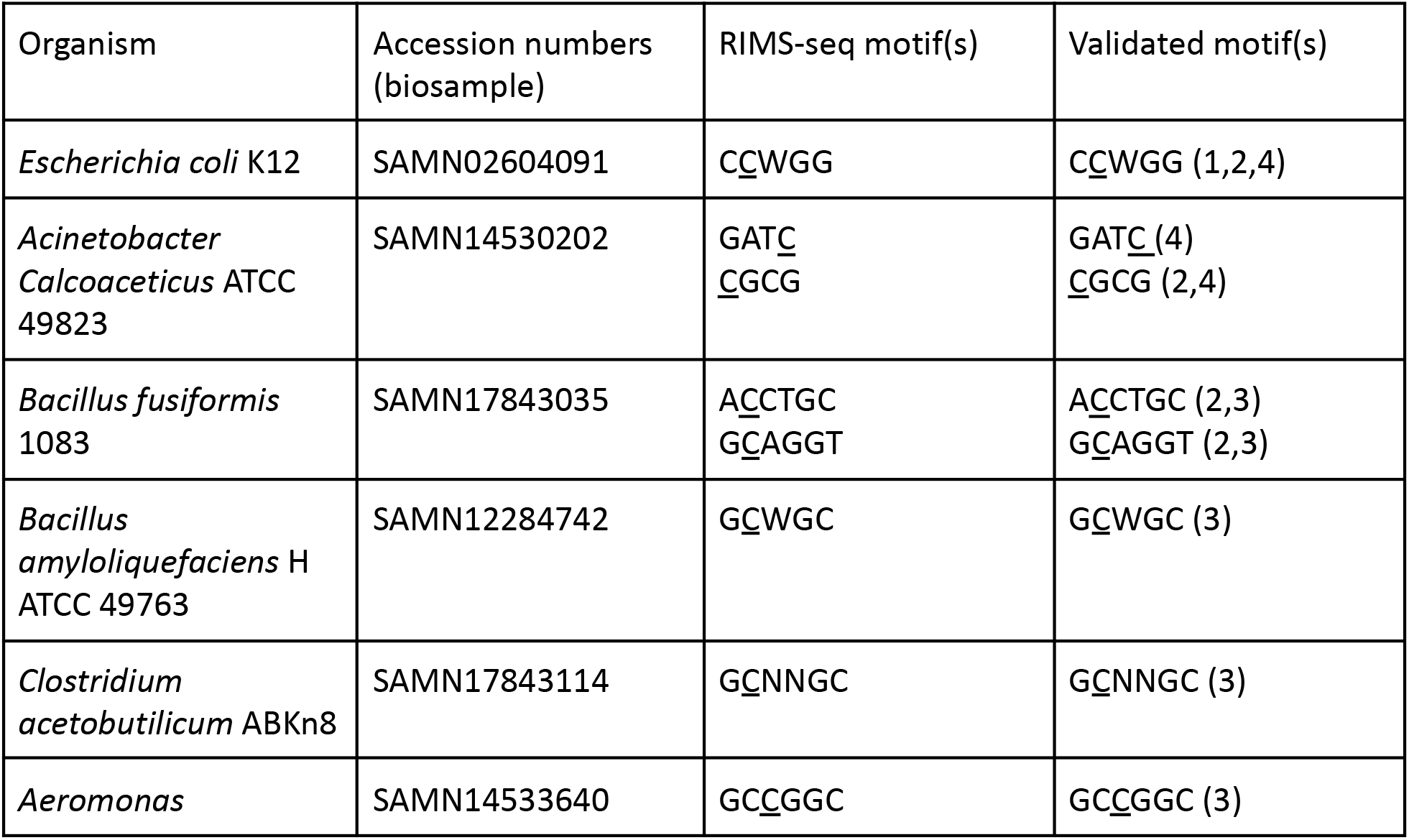
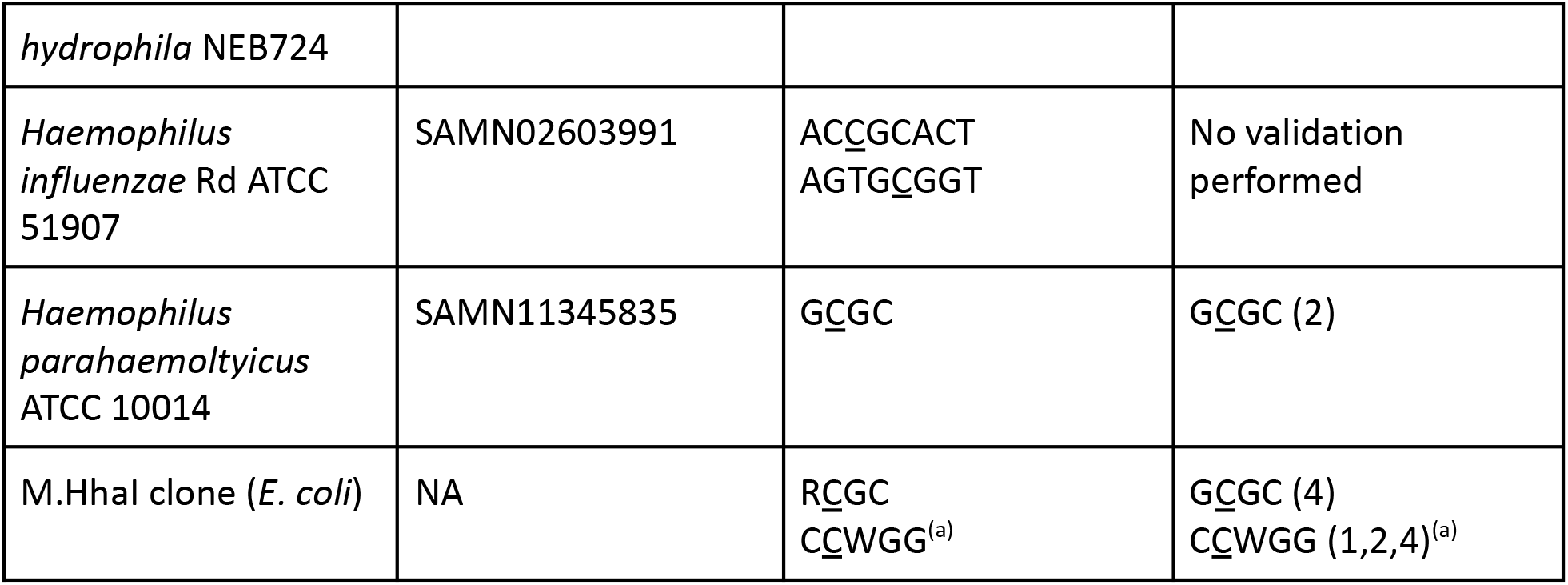
Methylases specificity obtained using RIMS-seq and validated using different methods. The method is indicated by a number next to the motif. : Evidence for the validated motifs are (1) Bisulfite-seq (material and Methods), (2) REBASE (Roberts et al., 2015) (3) EM-seq (material and method) (4) MFRE-seq (Anton et al., 2021). (a) The E. coli strain used is Dcm+, resulting in the discovery of both the Dcm (C<u>C</u>WGG) and M.HhaI motifs (G<u>C</u>GC). RIMS-seq discovered R<u>C</u>GC instead of G<u>C</u>GC motif (see text for explanation)

#### RIMS-seq identifies multiple methylase specificities *de novo* within a single microorganism

To assess whether RIMS-seq is able to identify methylase specificity in strains expressing multiple methylases, we repeated the same procedure on a strain of *Acinetobacter calcoaceticus* ATCC 49823 expressing two m5C methylases with known specificities (Roberts et al., 2015). RIMS-seq identifies <u>C</u>GCG (p-value =2.33e-174) and GAT<u>C</u> (p-value =3.02e-1308) (**Table 1**) both motifs have been confirmed by MFRE-seq (Anton et al., 2021). Thus, RIMS-seq is able to *de novo* identify methylase specificities in bacteria expressing multiple methylases.

#### RIMS-seq can be applied for genome sequencing and m5C profiling in bacteria without a reference genome

We investigated whether RIMS-seq can be used to identify methylase specificities of uncharacterized bacteria for which a reference genome is unavailable. More specifically we evaluated if the reads generated using RIMS-seq can be used for both identifying methylase specificities and generating an assembly of comparable quality to DNA-seq.

For this, we performed RIMS-seq on *A. calcoaceticus* ATCC 49823 genomic DNA as described above as well as a control DNA-seq experiment for which the alkaline treatment was replaced by 3 hours incubation in TE (DNA-seq(+3H)). We compared the *de novo* assembly obtained from the reads generated by the DNA-seq(+3H) and the *de novo* assembly obtained from the reads generated by RIMS-seq (see **Material and Methods**). In brief, the alkaline treatment did not alter the important metrics for assembly quality such as the rate of mismatches and N50 demonstrating that the elevated C to T variant rate at methylated sites is not high enough to cause assembly errors (**Figure 3C**).

We then proceeded to map the RIMS-seq reads to the assembly and motifs were identified using the RIMS-seq *de novo* motif discovery pipeline. As expected, the same motifs found when mapping to the reference genome are also found in the *A. calcoaceticus de novo* assembly with similar significance (GAT<u>C</u> (p-value=1.44e-1255) and <u>C</u>GCG (p-value=8.6e-228) (**Figure 3B**). These motifs correspond to the methylase specificities expected in this strain indicating that RIMS-seq can be applied for genome sequencing and assembly of any bacterium without the need for a reference genome.

#### RIMS-seq can be complemented with SMRT sequencing to obtain a comprehensive overview of methylase specificities

RIMS-seq performed in parallel with SMRT sequencing has the advantage of comprehensively identifying all methylase specificities (m5C, m4C and m6A methylations) and results in an assembly of higher quality than with short reads illumina data. We applied this hybrid approach to *Acinetobacter calcoaceticus* ATCC 49823 for which a SMRT sequencing and assembly had been done previously (Roberts et al., 2015). RIMS-seq was performed as described above and the reads were mapped to the genome assembly obtained from SMRT-sequencing. We again found the two m5C motifs : <u>C</u>GCG (p-value = 1.84e-1535) and GAT<u>C</u> (p-value = 4.93e-6856) from the RIMS-seq data in addition to the 13 m6A motifs described previously using SMRT sequencing (Roberts et al., 2015). This result demonstrates the advantage of such a hybrid approach in obtaining closed genomes with comprehensive epigenetic information.

### RIMS-seq can be applied to a variety of RM systems

Methylases targets are usually palindromic sequences between 4-8 nt, and a single bacterium often possesses several, distinct MTase activities (Vasu and Nagaraja, 2013). Next, we tested the general applicability of RIMS-seq and the *de novo* motif discovery pipeline using bacterial genomic DNA from our in-house collections of strains.

For some bacterial strains, the methylase recognition specificities have been previously experimentally characterized. In all of those strains, RIMS-seq confirms the specificities and identifies the methylated cytosine at base resolution (**Table 1**). We have tested the identification of 4-mers motifs such as GAT<u>C</u>, <u>C</u>GCG (*Acinetobacter calcoaceticus*) and G<u>C</u>GC (*Haemophilus parahaemolyticus*) up to 8-mers motifs such as AC<u>C</u>GCACT and AGTG<u>C</u>GGT (*Haemophilus influenzae*). Motifs can be palindromic or non-palindromic (**Table 1 and Supplementary Table 3**). In the latter case, RIMS-seq defines non-palindromic motifs at strand resolution. For example, RIMS-seq identifies methylation at two non-palindromic motifs A<u>C</u>CTGC as well as its reverse complement G<u>C</u>AGGT in the *Bacillus fusiformis* strain (**Table 1**).

A number of RM systems have been expressed in other hosts such as *E. coli* for biotechnological applications. For the methylase M.HhaI recognizing G<u>C</u>GC (Roberts et al., 2015), we performed RIMS-seq and a control DNA-seq(+3H) on both the native strain (*Haemophilus parahaemolyticus* ATCC 10014) and in *E. coli* K12 expressing the recombinant version of M.HhaI. Interestingly, we found that the *de novo* RIMS-seq analysis algorithm identifies R<u>C</u>GC (with R being either A or G) for the recombinant strain and G<u>C</u>GC for the native strain (**Figure 4A**). Conversely, no notable elevation of C to T read variants are observed for the native strain (**Figure 4B**), confirming the *de novo* motif discovery results from the analysis pipeline. Collectively, these results suggest that the recombinant methylase shows star activity, notably in the context of A<u>C</u>GC, that is not found in the native strain. We hypothesize that the star activity is the result of the over-expression of the methylase in *E. coli* K12. Interestingly, A<u>C</u>GC is not a palindrome motif and consequently the star activity results in hemi-methylation of the A<u>C</u>GC sites and not the GCGT motif.

**Figure 4 :**
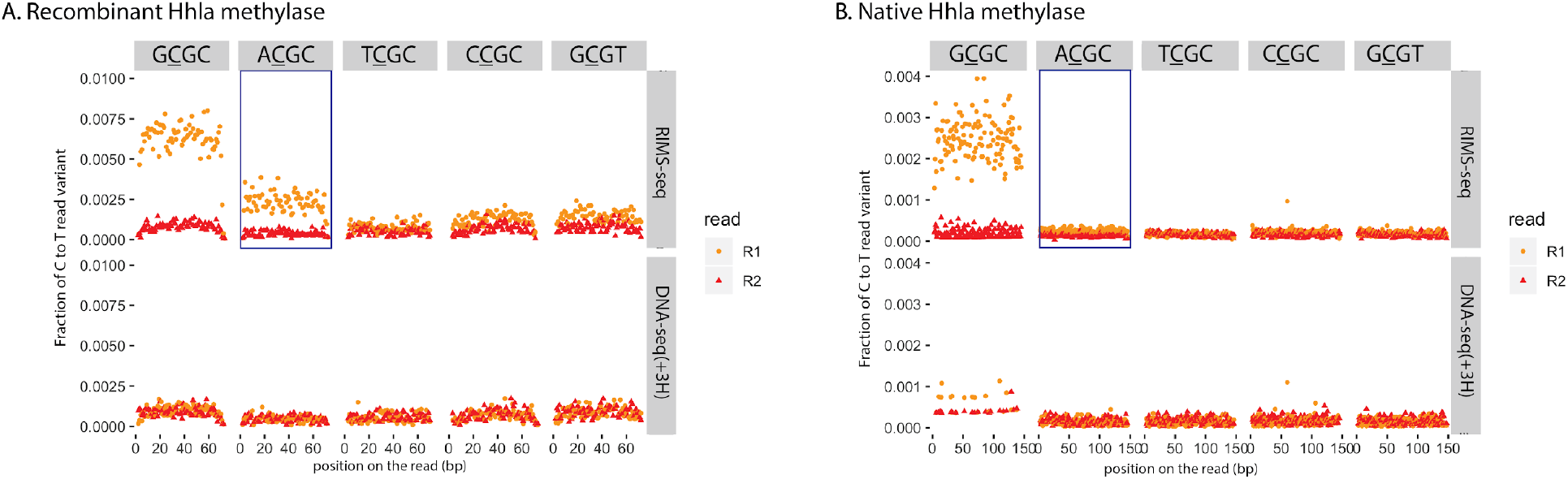
C to T error profile in G<u>C</u>GC (canonical recognition site), A<u>C</u>GC, T<u>C</u>GC, C<u>C</u>GC and G<u>C</u>GT. in R1 reads (orange) and R2 reads (red) for RIMS-seq (upper panel) and DNA-seq(+3H) (lower panel) *A.* Recombinant HhaI methylase expressed in *E.coli B.* Native HhaI methylase expressed in *Haemophilus parahaemolyticus*. Elevation of C to T in the R1 read variant can be observed in the context of G<u>C</u>GC for both the recombinant and native HhaI genomic DNA and in the context of A<u>C</u>GC only for DNA from the recombinant but not the native HhaI.

### RIMS-seq can be applied to microbial communities

We assessed whether RIMS-seq can be applied to mixed microbial communities using synthetic gut and skin microbiomes from ATCC containing 12 and 6 bacterial species, respectively. We also complemented the RIMS-seq experiment with the control experiment DNA-seq(+3H) and a bisulfite treatment to validate the RIMS-seq findings. Reads were mapped to their respective microbiome reference genomes (**Methods**). For the gut microbiome we found a mapping rate (properly paired only) of 95.79%, 95.77% and 66.2% for RIMS-seq, DNA-seq and bisulfite-seq respectively. Concerning the skin microbiome, 85.89%, 85.35% and 54.9% of reads were mapped for RIMS-seq, DNA-seq and bisulfite-seq respectively. The low mapping rate for bisulfite-seq is a known challenge as the reduction of the alphabet to A, G, T generates ambiguous mapping (Grehl et al., 2020).

To use RIMS-seq as an equivalent to DNA-seq for mixed community applications, RIMS-seq should produce sequencing quality metrics that are similar to standard DNA-seq, especially on the estimation of species relative abundances. We therefore compared RIMS-seq sequencing performances with DNA-seq(+3H) and bisulfite sequencing. We found that bisulfite sequencing elevates abundances of AT-rich species such as *Clostridioides difficile* (71% AT), *Enterococcus faecalis* (63% AT) and *Fusobacterium nucleatum* (73% AT) (**Figure 5A, Supplementary Figure 4**). For example, bisulfite sequencing over-estimated the presence of *Clostridioides difficile* by a factor of 2.65 and *Staphylococcus epidermidis* by a factor of 3.9 relative to DNA-seq. This over-estimation of an AT rich genome by bisulfite is a known bias of bisulfite sequencing and relates to damage at cytosine bases (Olova et al., 2018). Conversely, we found that the species abundances are similar between DNA-seq(+3H) and RIMS-seq (abundance ratios between 0.8 and 1.2) indicating that RIMS-seq can be used to quantitatively estimate microbial composition.

**Figure 5 :**
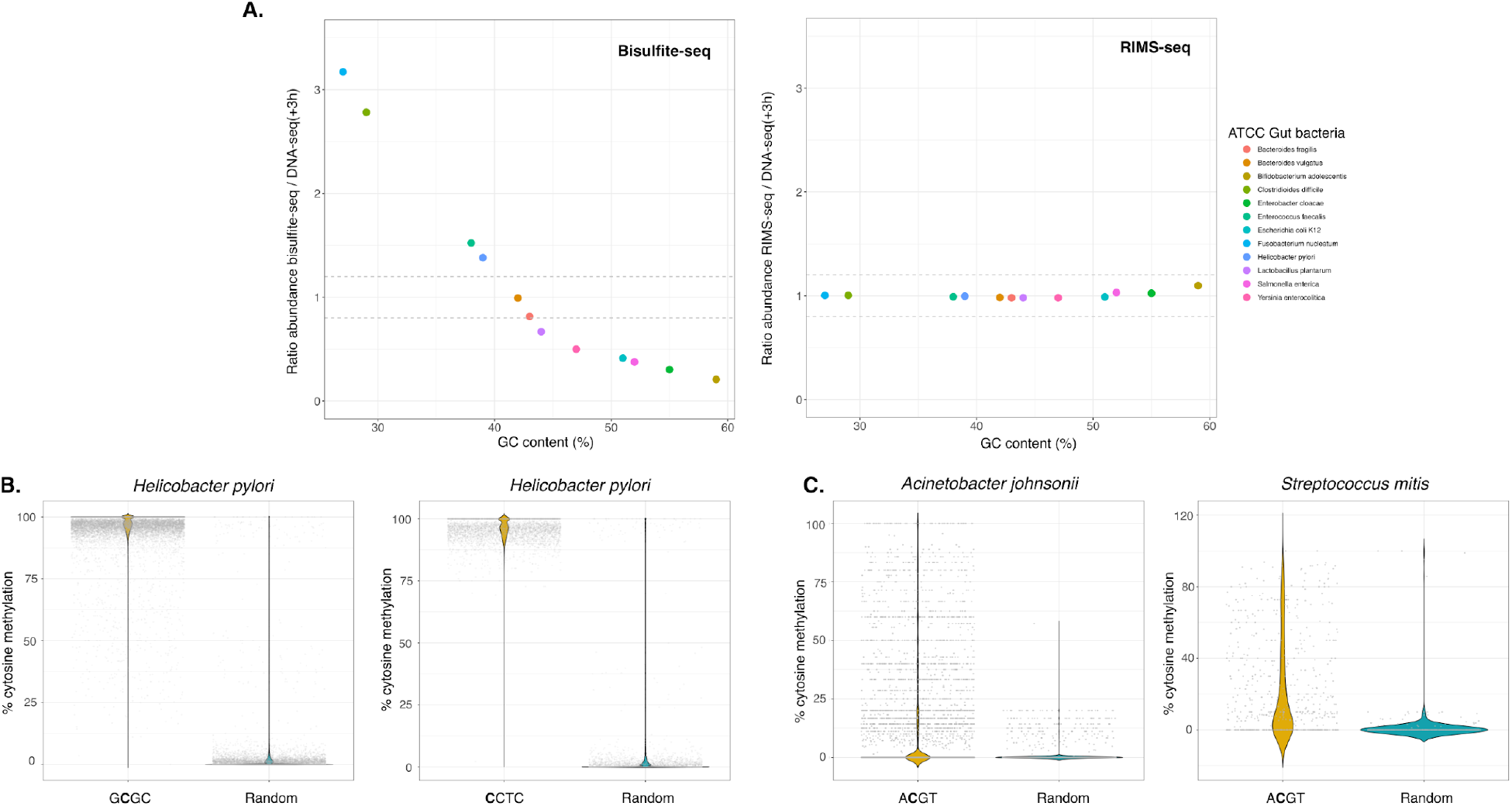
**A.** Bacterial abundance in the ATCC gut microbiome calculated from bisulfite-seq data (left) and RIMS-seq (Right) normalized to DNAseq(+3H). The normalized abundance is plotted relative to the GC content of each bacterium. **B.** Methylation level in *Helicobacter pylori* (ATCC gut microbiome) calculated from bisulfite-seq data. The methylation level was calculated for cytosine positions in the context of G<u>C</u>GC (yellow) and randomly selected positions in other contexts (blue). **C.** Methylation levels in *Acinetobacter johnsonii (left)* and *Streptococcus mitis (right)* (ATCC skin microbiome). The methylation level was calculated for cytosine positions in the context of A<u>C</u>GT (yellow) and randomly selected positions in other contexts (blue). These bisulfite-seq data suggest some sites are methylated in the context of A<u>C</u>GT, but they are not fully methylated.

#### RIMS-seq identifies known and novel methylase specificities in synthetic microbial communities

Overall, we found motifs for 6 out of the 12 gut microbiome species and 5 out of the 6 skin microbiome species (**Supplementary Table 3**). The motifs range from 4 to 8 nucleotides long and 70% are palindromic. Interestingly, we found an unknown palindromic motif GG<u>C</u>SGCC (with S being either C or G) from *Micrococcus luteus* (NC_012803.1) in the skin community. To our knowledge, this is the first time this 7nt motif is identified, showing the potential of RIMS-seq to identify new methylase specificities.

We validated the results obtained with RIMS-seq using bisulfite sequencing. RIMS-seq identified 2 motifs in *Helicobacter pylori* from the ATCC synthetic gut microbiome: G<u>C</u>GC as well as an additional non-palindromic motif <u>C</u>CTC that was identified by the bisulfite analysis pipeline as <u>C</u>YTC with Y being either C or T. The <u>C</u>CTC motif is very common in *Helicobacter pylori*s species, it has been described to be modified at m5C on one strand, while modified at m6A on the other strand (Roberts et al., 2015). In order to confirm the RIMS-seq motif, we investigated the bisulfite-seq data and compared the methylation level in cytosines present either in the <u>C</u>CTC context versus cytosines in any other context. We see a methylation level above 90% at the cytosines in the <u>C</u>CTC context confirming the existence of this methylated motif in *Helicobacter pylori* (**Figure 5B**). Interestingly, m4C methylation in *Helicobacter pylori* has been shown to also occur at T<u>C</u>TTC (Vitkute et al., 2001), resulting in the composite motif CYTC (T<u>C</u>TTC and N<u>C</u>CTC) found in the bisulfite data. Contrary to bisulfite, RIMS-seq does not identify m4C methylation (Vilkaitis and Klimasauskas, 1999).

Also interestingly, bisulfite-seq results indicate that the A<u>C</u>GT motif in *Acinetobacter johnsonii* and *Streptococcus mitis* from the ATCC synthetic skin microbiome are not fully methylated (**Figure 5C**). Most of the sites in *Acinetobacter johnsonii* show a methylation of about 10% while in *Streptococcus mitis*, the average methylation per site is 23%. These results highlight that despite the low methylation levels, RIMS-seq is able to detect the A<u>C</u>GT motif at high significance (p-value < 1e-100).

## Discussion

In this study, we developed RIMS-seq, a sequencing method to simultaneously obtain high quality genomic sequence and discover m5C methylase specificity(ies) in bacteria using a single library preparation. The simplicity of the procedure makes RIMS-seq a cost effective and time saving method with only an additional 3h sodium hydroxide incubation and an additional column-based cleaning step. Theoretically, the cleaning step can be avoided if a small volume of the library is used for the amplification step, but we have not tested this procedure.

Due to the limited deamination rate, RIMS-seq is equivalent to short read DNA-seq in terms of sequencing quality. Sequencing QC metrics such as coverage, GC content and mapping rate are similar for RIMS-seq and DNA-seq. Thus, RIMS-seq can be used for applications such as, but not limited to, shotgun sequencing, genome assembly and estimation of species composition of complex microbial communities. This dual aspect of RIMS-seq is analogous to SMRT sequencing for which methylation is inferred from the IPD ratio. We showed that both PacBio and RIMS-seq can be complementary with the ability to obtain a complete methylome: m6A and m4C methylase specificities can be obtained from SMRT sequencing while m5C methylase specificity can be obtained from RIMS-seq. Combining both sequencing technologies also allows for a hybrid assembly strategy resulting in closed reference genomes of high sequencing accuracy.

We applied RIMS-seq to several bacteria and identified a variety of methylation motifs, ranging from 4 to 8nt long, palindromic and non-palindromic. Some of these motifs were identified for the first time, demonstrating the potential of the technology to discover new methylase specificities, from known as well as from unknown genomes. We also validated that RIMS-seq can identify multiple methylase specificities from a synthetic microbial community and estimate species abundances. However, RIMS-seq has caveats similar to metagenomics sequencing when applied to study natural microbial communities. Closely related species are likely to co-exist and assigning the motif to the correct species can be challenging. Furthermore, single nucleotide polymorphisms may confound the identification of the C to T deamination increasing the background noise for the detection of motifs. Finally, species in microbiomes are unevenly represented which can cause RIMS-seq to identify motifs only in the most abundant species.

Because RIMS-seq is based on a limited deamination, it requires the combined signal over many reads to be large enough to effectively identify methylase specificity. For the vast majority of the methylases in RM systems, methylation is present at a sufficient number of sites across the genome for RIMS-seq to determine their specificities. Nonetheless, bacterial methylases can be involved in other processes such as, but not limited to, DNA mismatch repair (Modrich and Lahue, 1996), gene regulation (Casadesús and Low, 2006) and sporulation (Oliveira et al., 2020) and the recognition sites may not necessary be fully methylated. Partially methylated sites can be found using RIMS-seq but more analysis needs to be done to evaluate how pervasive methylation needs to be to provide a RIMS-seq signal. In other cases, methylated motifs are too specific or under purifying selection, resulting in just a handful of sites in the genome. In these cases, RIMS-seq signals can only be obtained with enough read coverage to compensate for the scarcity of those sites. While the methylase specificities are of great interest in bacteria due to their diversity in recognition sequences, applying RIMS-seq to humans would lead to the identification of the already well-described CpG context. In this case, other technologies such as EM-seq or bisulfite-seq are more appropriate as they enable the precise genomic location to be obtained.

In summary, RIMS-seq is a new technology allowing the simultaneous investigation of both the genomic sequence and the methylation in prokaryotes. Because this technique is easy to implement and shows similar sequencing metrics to DNA-seq, RIMS-seq has the potential to substitute DNA-seq for microbial studies.

## Material and Methods

### Samples and genomic DNA collection

Skin microbiome genomic DNA (ATCC^®^ MSA-1005) and gut microbiome genomic DNA (ATCC^®^ MSA-1006) were obtained from ATCC. *E. coli* BL21 genomic DNA was extracted from a culture of *E. coli* BL21 DE3 cells (C2527, New England Biolabs) using the DNEasy Blood and Tissue kit (69504, Qiagen). *E. coli* K12 MG1655 genomic DNA was extracted from a cell culture using the DNEasy Blood and Tissue kit (69504, Qiagen). All the other gDNA from the bacteria presented in **Table 1** were isolated using the Monarch genomic DNA purification kit (T3010S, New England Biolabs). Xp12 phage genomic DNA was obtained from Peter Weigele and Yian-Jiun Lee at New England Biolabs.

### RIMS-seq library preparation

100ng of gDNA was sonicated in 1X TE buffer using the Covaris S2 (Covaris) with the standard protocol for 50μL and 200bp insert size.

The subsequent fragmented gDNA was used as the starting input for the NEBNext Ultra II library prep kit for Illumina (E7645, New England Biolabs) following the manufacturer’s recommendations until the USER treatment step. The regular unmethylated loop-shaped adapter was used for ligation. After the USER treatment (step included), the samples were subjected to heat alkaline deamination: 1M NaOH pH 13 was added to a final concentration of 0.1M and the reactions were placed in a thermocycler at 60°C for 3h. Then, the samples were immediately cooled down on ice and 1M of acetic acid was added to a final concentration of 0.1M in order to neutralize the reactions.

The neutralized reactions were cleaned up using the Zymo oligo clean and concentrator kit (D4060 Zymo Research) and the DNA was eluted in 20μL of 0.1X TE.

PCR amplification of the samples was done following NEBNext Ultra II library prep kit for Illumina protocol (ER7645, New England Biolabs) and the NEBNext^®^ Multiplex Oligos for Illumina^®^(E7337A, New England Biolabs). The number of PCR cycles was tested and optimized for each sample following the standard procedure for library preparation. PCR reactions were cleaned up using 0.9X NEBNext Sample purification beads (E7137AA, New England Biolabs) and eluted in 25μL of 0.1X TE. All the libraries were evaluated on a TapeStation High sensitivity DNA1000 (Agilent Technologies) and paired-end sequenced on Illumina.

### Bisulfite-seq library preparation

1% of lambda phage gDNA (D1221, Promega) was spiked-into 300ng gDNA to use as an unmethylated internal control. The samples were sonicated in 1X TE buffer using the Covaris S2 (Covaris) with the standard protocol for 50μL and 200bp insert size.

The subsequent fragmented gDNA was used as the starting input for the NEBNext Ultra II library prep kit for Illumina (E7645, New England Biolabs) following the manufacturer’s recommendations until the USER treatment step. The methylated loop-shaped adapter was used for ligation. After USER, a 0.6X clean-up was performed using the NEBNext Sample purification beads (E7137AA, New England Biolabs) and eluted in 20μL of 0.1X TE. A TapeStation High Sensitivity DNA1000 was used to assess the quality of the library before subsequent bisulfite treatment. The Zymo EZ DNA Methylation-Gold Kit (D5005, Zymo Research) was used for bisulfite treatment, following the manufacturer’s suggestions.

PCR amplification of the samples was done following the suggestions from NEBNext Ultra II library prep kit for Illumina (ER7645, New England Biolabs), using the NEBNext^®^ Multiplex Oligos for Illumina^®^(E7337A, New England Biolabs) and NEBNext^®^ Q5U^®^ Master Mix (M0597, New England Biolabs).

The number of PCR cycles was tested and optimized for each sample. The PCR reactions were cleaned up using 0.9X NEBNext Sample purification beads (E7137AA, New England Biolabs) and eluted in 25μL of 0.1X TE. All the libraries were screened on a TapeStation High sensitivity DNA1000 (Agilent Technologies) and paired-end sequenced on Illumina.

### RIMS-seq data analysis

Paired-end reads were trimmed using Trim Galore 0.6.3 (option --trim1). The *Acinetobacter calcoaceticus* ATCC 49823 data have been trimmed using Trim Galore version 0.6.3 instead and downsampled to 1 million reads. Reads were mapped to the appropriate genome using BWA mem with the paired-end mode (version 0.7.5a-r418 and version 0.7.17-r1188 for the *Acinetobacter calcoaceticus*). When using an assembled genome directly from RIMS-seq data, trimmed RIMS-seq reads were assembled using SPAdes (SPAdes-3.13.0 (Nurk et al., 2013)default parameters). Reads were split according to the read origin (Read 1 or Read 2) using samtools (version 1.8) with -f 64 (for Read 1) and -f 128 (for Read 2) and samtools mpileup (version 1.8) was run on the split read files with the following parameters : -O -s -q 10 -Q 0. For *Acinetobacter calcoaceticus*, the unmapped reads, reads without a mapped mate and the non primary alignments were filtered out using the flags -F 12 and -F 256.

### De-novo identification of motifs using RIMS-seq

Programs and a detailed manual for the de-novo identification of motifs in RIMS-seq are available on github (https://github.com/Ettwiller/RIMS-seq/). Using the mpileup files, positions and 14bp flanking regions in the genome for which a high quality (base quality score >= 35) C to T in R1 or a G to A in R2 was observed were extracted for the foreground. Positions and 14bp flanking regions for which a high quality (base quality score >= 35) G to A in R1 or a C to T in R2 was observed were extracted for the background. C to T or G to A in the first position of reads were ignored. If the percentage of C to T or G to A are above 5% for at least 5 reads at any given position, the position was ignored (to avoid considering positions containing true variants). Motifs that are found significantly enriched (p-value < 1e-100) in the foreground sequences compared to background sequences were found using mosdi pipeline mosdi-discovery with the following parameters : ‘*mosdi-discovery -v discovery -q x -i -T 1e-100 -M 8,1,0,4 8 occ-count’* using the foreground sequences with x being the output of the following command : ‘*mosdi-utils count-qgrams -A “dna”* ‘ using the background sequences.

To identify additional motifs, the most significant motif found using *mosdi-discovery* is removed from the foreground and background sequences using the following parameter : ‘*mosdi-utils cut-out-motif -M X’* and the motif discovery process is repeated until no motif can be found.

### Sequence logo generation

Using the mpileup files, positions in the genome for which a high quality (base quality score >= 35) C to T in R1 or a G to A in R2 was observed were extracted for the foreground using the get_motif_step1.pl program. Positions for which a high quality (base quality score >= 35) G to A in R1 or a C to T in R2 was observed were extracted for the background. The +/-7bp regions flanking those positions were used to run Two sample logo (Vacic et al., 2006). Parameters were set as t-test, p.value < 0.01.

### Bisulfite-seq data analysis

Reads were trimmed using Trim Galore 0.6.3 and mapped to the bisulfite-converted concatenated reference genomes of each respective synthetic microbiome using bismark 0.22.2 with default parameters. PCR duplicates were removed using deduplicate_bismark and methylation information extracted using bismark_methylation_extractor using default parameters. For the microbiome, the bismark_methylation_extractor with --split_by_chromosome option was used to output one methylation report per bacterium. The motif identification was done as previously described in (Anton et al. 2021).

### EM-seq

EM-seq was performed according to the standard protocol (NEB E7120S) Motif identification was done as previously described in (Anton et al., 2021).

### Analysis and abundance estimation in synthetic microbiomes

RIMS-seq, DNA-seq and Bisulfite-seq were performed on the synthetic gut and skin microbiome as described. Reads derived from RIMS-seq, DNA-seq and Bisulfite-seq were mapped as described to a ‘meta-genome’ composed of the reference genomes of all the bacteria included in the corresponding synthetic community (see Supplementary Table 3 for detailed compositions). Mapped reads were split according to each bacterium and RIMS-seq or bisulfite analysis pipelines were run on individual genomes as described above. Abundance was estimated using the number of mapped reads per bacteria and normalized to the total number of mapped reads. Normalized species abundances in RIMS-seq and Bisulfite-seq were compared to the normalized species abundances in DNA-seq.

### Quality control of the data

The insert size for each downsampled filtered bam file was calculated using Picard version 2.20.8 using the default parameters and the option CollectInsertSizeMetrics (“Picard Toolkit.” 2019. Broad Institute, GitHub Repository. http://broadinstitute.github.io/picard/; Broad Institute).

The GC bias for each downsampled filtered bam file was calculated and plotted using Picard version 2.20.8 using the default parameters and the option CollectGcBiasMetrics.

### Xp12 genome assembly

Reads were downsampled to a 30X coverage using seqtk 1.3.106, trimmed using trimgalore 0.6.5 and assembled using Spades 3.14.1 with the --isolate option. Assembly quality was assessed using Quast 5.0.2. Reads used for assembly were then mapped back to the assembly using BWA mem 0.7.17 and mapping statistics were generated using samtools flagstat 1.10.2

### Xp12 sequencing performance assessment

Reads were trimmed using trimgalore 0.6.5 and mapped to the Xp12 reference genome using BWA mem 0.7.17. Insert size and GC bias were assessed using Picard Toolkit and genome coverage using Qualimap 2.1.1.

## Code and data availability

The data have been deposited with links to BioProject accession number PRJNA706563 in the NCBI BioProject database (https://www.ncbi.nlm.nih.gov/bioproject/).

Custom-built bioinformatics pipelines to analyse sequencing reads from RIMS-seq are available at https://github.com/Ettwiller/RIMS-seq/

## Acknowledgments

We thank Peter Weigele and Yian-Jiun Lee from New England Biolabs for the Xp12 genomic DNA and genomic sequence. We thank Ivan Correa and Nan Dai for their assistance with LC-MS and Ira Schildkraut for his help with methylase specificities.

## Competing interests

CB, YCL, AF, BPA, LC, TCE, RR and LE are or were employees of New England Biolabs Inc. a manufacturer of restriction enzymes and molecular reagents.

## Notes

https://github.com/Ettwiller/RIMS-seq/

