## Supplementary Materials for "Rapid Identification of Methylase Specificity (RIMS-seq) jointly identifies methylated motifs and generates shotgun sequencing of bacterial genomes"

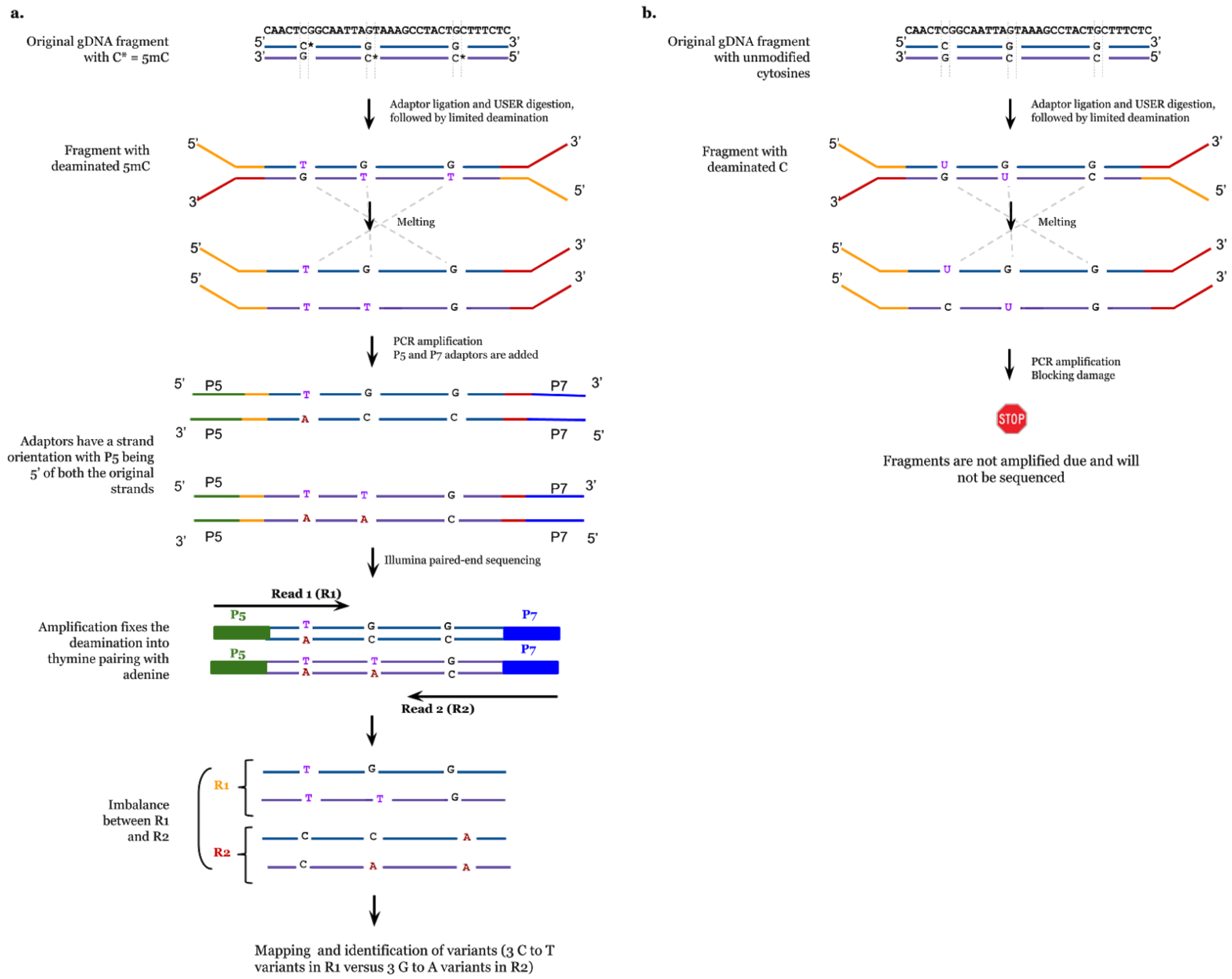


**Supplementary Figure 1 : a.** Scheme highlighting the imbalance between C to T read variants in the first paired-end read (R1) compared to the second paired-end read (R2).
Following fragmentation of gDNA fragments containing 5mC, loop-shaped adaptors are ligated and linearized with USER (see Methods). The fragments are then subjected to a limited deamination, leading to 5mC to T conversion. The illumina sequencing adaptors P5 and P7 are added during PCR amplification, enabling subsequent strand-oriented sequencing : P5 adaptor is added to the 5'end of the original fragment while P7 is added to the 3'end of the original fragment. During illumina sequencing, sequencing from the P5 adaptor (Read 1) corresponds to the original strand; while sequencing from the the P7 adaptor, corresponds to the reverse complement of the original strand. Because the deamination affects only one base of a pair (5mC), this leads to an increase of C to T read variants in Read 1 (original strand), whereas Read

2 shows an increase of the reverse complement of C to T : an increase of G to A read variants. This results in a global imbalance of C to T read variants in Read 1 compared to Read 2.
**b.** Scheme highlighting the effect of the deamination of regular cytosines. Following fragmentation of gDNA fragments containing regular C, loop-shaped adaptors are ligated and linearized with USER. The fragments are then subjected to a limited deamination, leading to C to U conversion. This is a blocking damage for the polymerase that cannot amplify fragments containing Us. Thus, P5 and P7 illumina adaptors are not added and the fragments will not be sequenced.

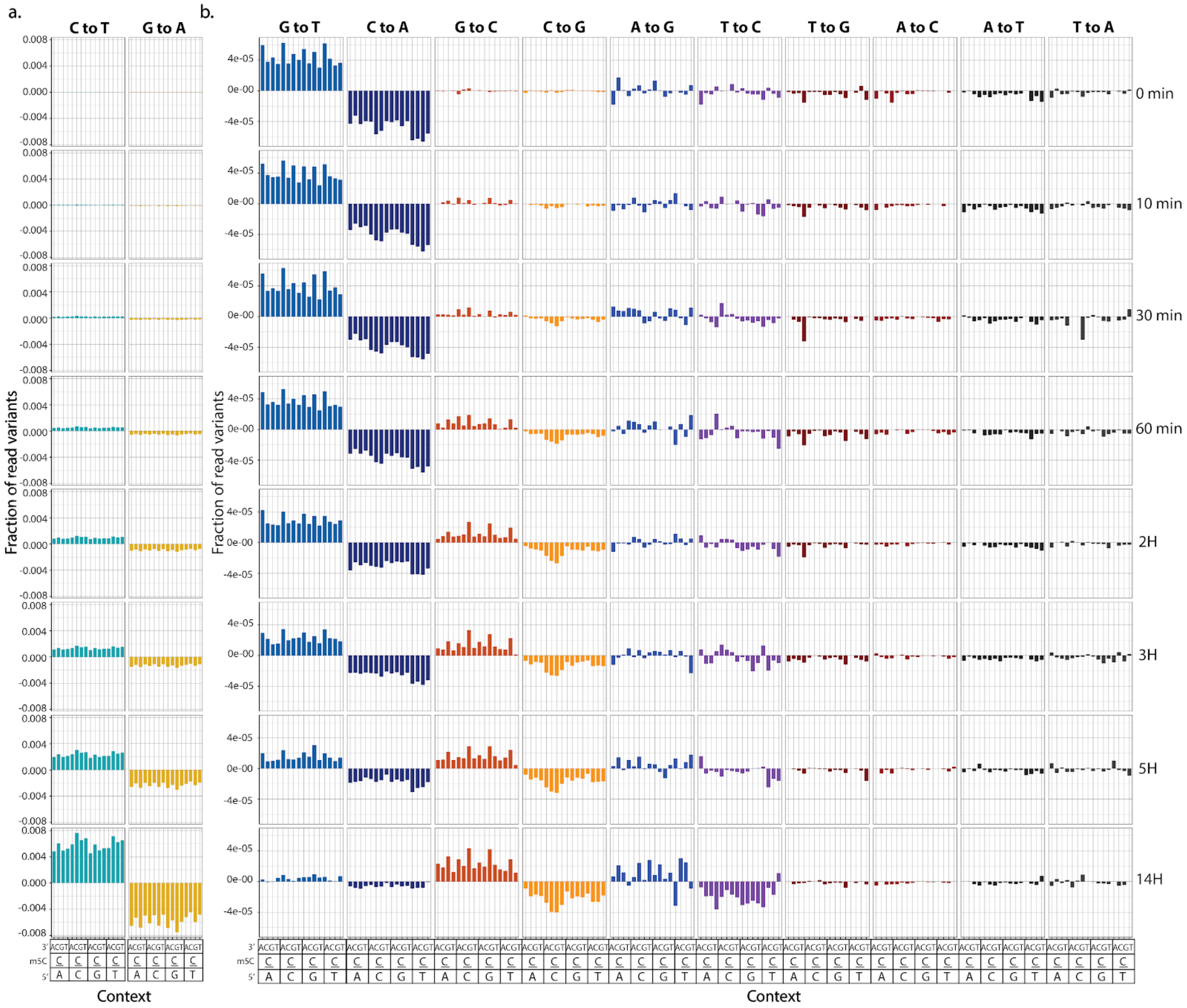


**Supplementary Figure 2** Profile of read variants in a fully methylated genome (XP12) after RIMS-seq. **a.** Excess of read variants for control (t=0) and various heat alkaline treatment times (t= 10, 30 minutes, 1,2,3,5 and 14 hours) for C to T and G to A. **b.** Same as a. for all the other substitutions. Time represents the heat-alkaline incubation time (in minutes or hours). C to T variants are increasing over time. G to C shows a moderate increase over time. Conversely G to T variants are decreasing over time. Note that the scale between **a.** and **b.** is different and the C to T excess in R1 is up to a 100 fold greater than for the other substitutions.

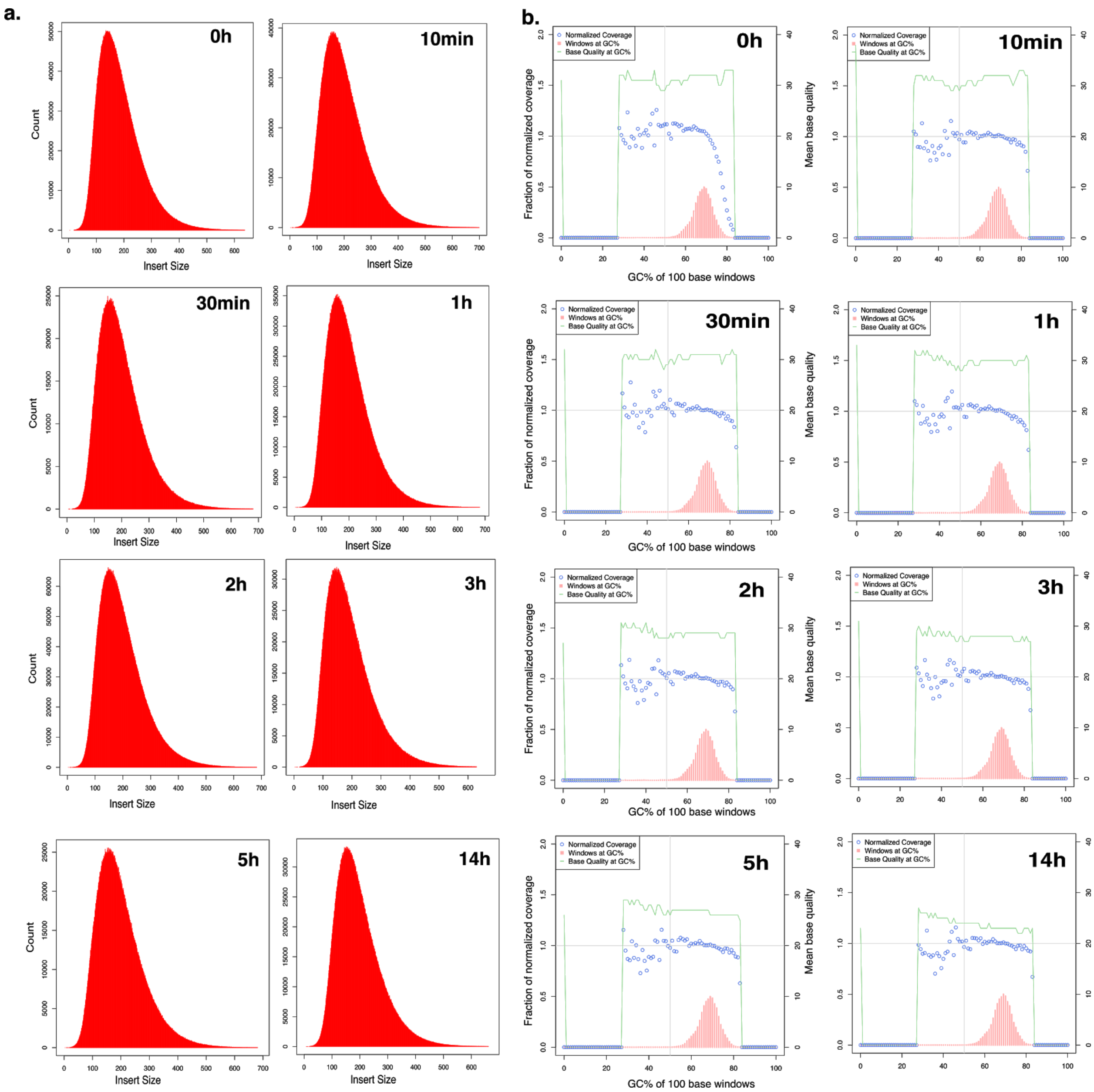


**Supplementary Figure 3: Quality control of the sequencing performances for Xp12 gDNA**

**a.** Insert size distribution (bp) for the control (0h) and different heat-alkaline treatment times. Input genomic DNA was sheared by Covaris treatment (200 bp target) and no further size selection was applied.

**b**. GC bias for the control (0h) and different heat-alkaline treatment times.

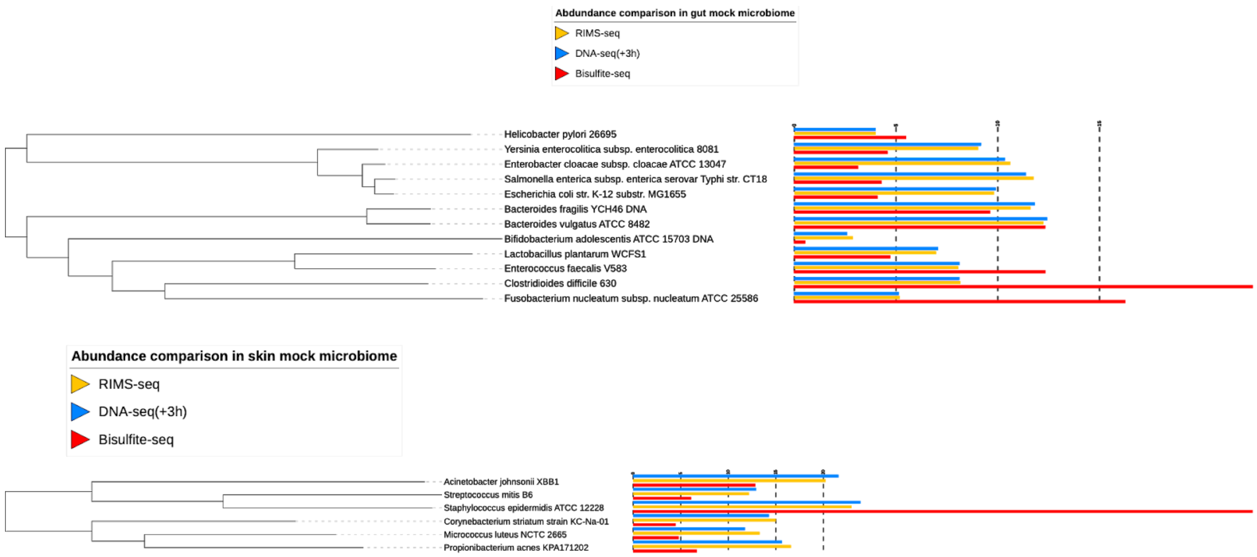


**Supplementary Figure 4**: Phylogenetic tree and barplot showing the relative abundance of each species in the gut (top) and skin (bottom) microbiome for RIMS-seq (yellow), DNA-seq(+3H, blue) and bisulfite sequencing (red). Relative abundance is calculated using the number of reads mapping to each species normalized to  the total number of mapped reads.

| Alakline-heat treatment time | 0h | 10min | 30min | 1h | 2h | 3h | 5h | 14h |
|---|---|---|---|---|---|---|---|---|
| **Statistics without reference genome** | | | | | | | | |
| nb contigs | 1 | 1 | 1 | 1 | 1 | 1 | 1 | 1 |
| largest contig | 63783 | 63882 | 64181 | 63774 | 63873 | 63839 | 63873 | 63774 |
| **Statistics with reference genome** | | | | | | | | |
| largest alignment | 63783 | 63782 | 63774 | 63774 | 63285 | 63839 | 63773 | 63774 |
| total aligned length | 63783 | 63782 | 63774 | 63774 | 63285 | 63839 | 63773 | 63774 |
| GC % | 68.17 | 68.17 | 68.17 | 68.17 | 68.17 | 68.18 | 68.17 | 68.17 |
| N50 | 63783 | 63882 | 63774 | 63774 | 63873 | 63839 | 63873 | 63774 |
| Genome fraction (%) | 99.239 | 99.238 | 99.225 | 99.225 | 99.224 | 99.241 | 99.224 | 99.225 |
| % reads mapping back to assembly | 99.75 | 99.73 | 99.8 | 99.76 | 99.82 | 99.78 | 99.75 | 99.84 |
| **Misassemblies** | | | | | | | | |
| nb misassemblies | 0 | 0 | 0 | 0 | 0 | 0 | 0 | 0 |
| misassembled contig length | 0 | 0 | 0 | 0 | 0 | 0 | 0 | 0 |
| local misassemblies | 1 | 1 | 1 | 1 | 1 | 1 | 1 | 1 |
| **Mismatches** | | | | | | | | |
| N's per 100kbp | 0 | 156.14 | 0 | 0 | 156.56 | 0 | 156.56 | 0 |
| nb mismatches per 100kbp | 3.14 | 3.14 | 1.57 | 1.57 | 3.14 | 4.7 | 1.57 | 3.14 |
| nb indels per 100kbp | 0 | 0 | 0 | 0 | 0 | 0 | 0 | 0 |

**Supplementary Table 1:** Xp12 assembly statistics (see material and methods)

| | dnaseq 3h | RIMS | reference |
|---|---|---|---|
| span (bp) | 3,540,609 | 3,515,857 | 3,543,981 |
| N (%) | 0.02 | 0.01 | 0.00 |
| GC (%) | 38.62 | 38.59 | 38.31 |
| AT (%) | 61.38 | 61.41 | 61.69 |
| scaffold count | 571 | 531 | 1 |
| longest scaffold (bp) | 317,976 | 318,149 | 3,543,981 |
| scaffold N50 length (bp) | 77,278 | 62,717 | 3,543,981 |
| scaffold N50 count | 14 | 15 | 1 |
| scaffold N90 length (bp) | 14,355 | 13,679 | 3,543,981 |
| scaffold N90 count | 52 | 59 | 1 |
| contig count | 588 | 540 | 1 |
| contig N50 length (bp) | 62,601 | 58,046 | 3,543,981 |
| contig N50 count | 15 | 16 | 1 |
| contig N90 length (bp) | 13,997 | 13,455 | 3,543,981 |
| contig N90 count | 57 | 63 | 1 |

**Supplementary Table 2 :** Assembly statistics for *Acinetobacter calcoaceticus* ATCC 49823 assemblies obtained using the sequences from the standard DNA-seq (+3H) and RIMS-seq, compared to the reference genome.

| Skin microbiome | RIMS-seq motif | Bisulfite-seq motif | p-value RIMS-seq | Chromosome accession | GC content (%) |
|---|---|---|---|---|---|
| *Micrococcus luteus* | GG**C**SGCC | GG**C**SGCC | 7.965455e-572 | NC_012803.1 | 73 |
| | **C**GSNNW | | 1.74E-154 | | |
| *Propionibacterium acnes* | CSNNNN**C**G | NA | 2.34E-132 | NC_006085.1 | 60 |
| *Corynebacterium striatum* | G**C**CGGC | G**C**CGGC | 1.37E-268 | NZ_CP021252.1 | 59 |
| | **C**NNYRNNG | | 4.72E-135 | | |
| *Acinetobacter johnsonii* | A**C**GT | A**C**GT | 1.19E-105 | NZ_CP010350.1 | 41 |
| | AT**C**NNGRC | | 3.12E-123 | | |
| *Streptococcus mitis* | G**C**NGC | G**C**NGC | 1.540920e-603 | NC_013853.1 | 40 |
| | A**C**GT | | 6.07E-108 | | |
| *Staphylococcus epidermidis* | NA | NA | NA | NC_004461.1 | 32 |
| **Gut microbiome** | **RIMS-seq motif** | **Bisulfite-seq motif** | **p-value RIMS-seq** | **Chromosome accession** | **GC content (%)** |
| *Bifidobacterium adolescentis* | GAT**C** | GAT**C** | 7.609400e-718 | NC_008618.1 | 59 |
| | C**C**NGG | C**C**NGG | 3.970975e-364 | | |
| *Enterobacter cloacae* | C**C**WGG | C**C**WGG | 3.129282e-1371 | NC_014121.1 | 55 |
| *Salmonella enterica* | C**C**WGG | C**C**WGG | 1.071714e-1331 | NC_003198.1 | 52 |
| *Escherichia coli K12* | C**C**WGG | C**C**WGG | 2.277395e-1220 | NC_000913.3 | 51 |
| *Yersinia enterocolitica* | C**C**WGG | C**C**WGG | 2.314551e-649 | NC_008800.1 | 47 |
| *Lactobacillus plantarum* | NA | NA | NA | NC_004567.2 | 44 |
| *Bacteroides fragilis* | NA | NA | NA | NC_006347.1 | 43 |
| *Bacteroides vulgatus* | NA | NA | NA | NC_009614.1 | 42 |
| *Helicobacter pylori* | G**C**GC | G**C**GC | 2.291291e-516 | NC_000915.1 | 39 |
| | **C**CTC | | 2.387073e-318 | | |
| *Enterococcus faecalis* | NA | NA | NA | NC_004668.1 | 38 |
| *Clostridioides difficile* | NA | NA | NA | NC_009089.1 | 29 |
| *Fusobacterium nucleatum* | NA | NA | NA | NC_003454.1 | 27 |

**Supplementary Table 3 :** methylases specificity of the synthetic ATCC microbiomes
